# Feedforward representation of face information in the fusiform face area revealed by VASO 7T layer fMRI

**DOI:** 10.1101/2025.11.04.686314

**Authors:** Kenshu Koiso, Kazuaki Akamatsu, Renzo Huber, Yoichi Miyawaki

## Abstract

Recognizing faces is a critical cognitive process in social interaction. Face recognition has been extensively studied with functional magnetic resonance imaging (fMRI), ranging from large coverage studies comprehensively examining the entire face pathway to high-resolution laminar-specific (layer) fMRI studies probing feedforward and feedback signals in early visual and face-selective areas during particular face processing. However, it is still unclear which brain region structures face information in the face pathway. To further elucidate this mechanism, we investigated fusiform face area (FFA) during passive face image presentation using a whole-brain vascular space occupancy (VASO) 7T layer fMRI open dataset combined with multi-voxel pattern analysis (MVPA). We analysed lateral occipital cortex (LOC) as a control region and BOLD data in the dataset as a control contrast. We trained multiple binary classification decoders with various image categories (e.g., face vs. others, house vs. others) and three cortical layer groups independently to assess information representation across cortical depths. Our results revealed that decoding accuracies in VASO data peaked in the middle layer of FFA, suggesting a feedforward signature specifically during face processing. This effect was not observed in LOC or in decoders for other image categories. These findings indicate that face information is structured prior to processing in the FFA, consistent with previous reports suggesting that face-related features are partially extracted before reaching higher-level face areas. Overall, this study highlights the potential of combining high-specificity VASO layer-fMRI with high-sensitivity MVPA to dissect cortical information flow.

## Introduction

Recognizing faces is a critical cognitive process in our social context. We can complete this process without being conscious of it at all. Many studies have investigated how and with what pathway we process face information in the visual system (Bernstein & Yovel, 2015;Grill-Spector et al., 2017; Kanwisher et al., 1997). Our understanding of face recognition has advanced remarkably in the last century, driven by the development of invasive and non-invasive neuroimaging methods (Sellal, 2022). In particular, some studies have highlighted the unique aspects and benefits of high-resolution functional magnetic resonance imaging (laminar-specific fMRI, or layer fMRI) in face areas (Carricarte et al., 2024; Dowdle et al., 2022). Dowdle et al. (2022) used ambiguous face images as stimuli with two independent tasks that control attention to the face images (face detection task and fixation detection task) and showed a feedback signature in fusiform face area (FFA) uniquely when taking the contrast of laminar signals of these two tasks. Carricarte et al. (2024) showed the distinction between feedforward and feedback signatures in FFA, presenting well-known face image stimuli and their mental imagery.

Although many studies have investigated face pathway and face processing locus, it is still unclear where the face information is structured. For static face perception, F. Nichols et al. (2010) showed that information about facial parts exists not only in FFA but also in an earlier area, occipital face area (OFA), which is believed as an area strongly related to processed face information using multivoxel pattern analysis (MVPA). They also suggested that complete face information is more pronouncedly represented in FFA. This evidence indicates that preselected partial face information goes into FFA and is processed in FFA. Namely, one could speculate to find a feedforward signature in FFA in face information flow during passive face image presentation. Yet, to our knowledge, no study has shown a feedforward signature in FFA during passive face image presentation.

Moreover, previous studies argued that layer-specific information in the human brain was dissociated by the combination of layer fMRI and multivoxel pattern analysis (MVPA) (Muckli et al., 2015). They depicted the deep-layer-driven feedback signature in a mental imaginary data. Hence, this approach uniquely shows high detectability of informative brain activity patterns even when we cannot see statistically significant univariate signals in noisy data.

MVPA is also thought to have a reduced bias to large draining veins in gradient echo blood oxygenation level dependent (GE-BOLD) contrast for layer fMRI (Moerel et al., 2018; Muckli et al., 2015; Vizioli et al., 2020). Thus, MVPA can help increase the spatial specificity of fMRI signals. However, it remains controversial if residual sensitivities to large draining veins exist.

Hence, it would be informative to compare the results of MVPA applied to GE-BOLD and non-BOLD vein-free contrast, such as vascular space occupancy (VASO), even if this contrast is known to have much less sensitivity compared to GE-BOLD contrast (Haenelt et al., 2023; Iamshchinina et al., 2021).

Investigating face areas with layer fMRI without venous bias is also challenging in terms of signal acquisition. Typically, we prefer to use ultra-high field (7 T or higher) MRI to mitigate the decrease of signal-to-noise ratio (SNR) accompanying high-resolution scans. However, B0-shimming is especially difficult in the bottom part of the brain where face areas are located. Hence, we are sacrificing a lot of signals by acquiring VASO signals in these brain areas. That is why combining VASO fMRI with MVPA can be particularly beneficial, as MVPA’s improved sensitivity can potentially compensate for VASO’s low SNR.

In this study, we used a whole-brain VASO layer fMRI dataset to address face information representation across cortical layers in the visual object category-selective areas (fusiform face area (FFA) and lateral occipital cortex (LOC)) with MVPA. For this purpose, we used an open dataset acquired in our previous study (Koiso et al., 2023). Although this dataset has only one participant observing naturalistic movies in the main session, it is the only open VASO dataset currently available for investigating the representation of face information across the cortical layers with high reliability. Using this dataset, we examined whether face information processing is done prior to FFA, namely, face information would be represented in FFA as a feedforward signature. Additionally, we aim to investigate whether the VASO layer fMRI can provide additional information to conventional BOLD layer fMRI.

## Methods

### Dataset

We used an open dataset of the whole-brain layer fMRI (Kenshu dataset (Koiso et al., 2023)) that contains BOLD and VASO signals simultaneously acquired at a 7T Siemens MAGNETOM scanner. 3D-EPI (Poser et al., 2013), slice selective slab-inversion (SS-SI) VASO (Huber et al., 2014) with TR_vol_ of 5.1-5.2 s, 0.8 mm^3^ isotropic spatial resolution was used. This dataset was acquired while a participant observed a collection of naturalistic movies lasting 15 min (Human Connectome Project (HCP) “movie1” (Mekete & Vu, 2017), 51 times across 10 experimental sessions with free fixation. This dataset also comes along with structural reference data from 3T at 0.8 mm^3^ isotropic. We used the preprocessed data (motion correction, BOLD correction (removing residual BOLD effect in VASO signal by dividing blood-nulled data by non-nulled data, see details in Huber et al., 2014), and registration between functional and structural images) posted on OpenNeuro (https://openneuro.org/datasets/ds003216). For further details, see the dataset paper (Koiso et al., 2023).

### Decoding procedure

#### Preprocessing

Preprocessing has been done in AFNI version 20.3.05 ‘Vespasian’. First, to improve tSNR, all 51 runs of VASO/BOLD functional data (blood-nulled and non-nulled data) were averaged. To avoid noise amplification by multiple BOLD corrections, we took the average before BOLD corrections in nulled data (https://layerfmri.com/baddi/). Then, the time series were detrended to account for hardware-related signal drifts across the relatively long runs using the third polynomial function.

#### Defining the label, region of interest, and layer group

Among all volumes in the averaged data, we defined volumes with target labels (such as “face” and “house”) as “target volumes” and those without target labels as “others volumes”, and performed binary classification analysis. We selected the target volumes according to HCP WordNet. Since there is a mismatch in temporal resolution between HCP WordNet definition and TR_vol_ (HCP WordNet definition is every second and our TR_vol_ is approximately 5 sec), we defined the target volume when objects with the target labels were presented for more than half the duration of their TR_vol_. We included all noun labels that corresponded to more than 10 TR_vol_ as the target volumes. 14 labels were used as target volumes (“building”, “car”, “face”, “fence”, “hand”, “hat”, “house”, “land”, “man”, “road”, “sky”, “text”, “tree”, and “vegetation”). Also, we randomly selected the same number of “others” volumes that did not include target labels in their TR_vol_ at all.

We defined the FFA and LOC voxels by using MNI functional parcellation (Glasser et al., 2016) registered in the native space of the Kenshu dataset. As mentioned in the dataset paper, ANTs (version 2.1.0) non-linear alignment was performed for the registration (check the dataset paper for more details). Furthermore, we visually inspected the image distortion of these areas.

Layerification was done in volume space. We first defined 11 layers from the manually corrected anatomical segmentation file by LayNii (version 2.2.0), “LN2_LAYERS”. After that, the 11 layers were grouped into three layer groups as described in the dataset paper (see Koiso et al., 2023 for more details).

#### Training and evaluation

Linear Support Vector Machines (SVM) (python 3.8.5, scikit-learn 1.0.2) were trained as decoders to predict target or “others” volumes from VASO and BOLD data in layer groups of each region of interest independently. To quantify the decoder’s performance, we performed leave-one-paired-volume-out cross-validation, in which one target volume and one “others” volume were set aside for testing, and the remaining volumes were used for training. We performed the “others” volume selection procedure 100 times and evaluated the prediction accuracies of the decoders to mitigate the potential bias caused by selecting a part of the data. Numbers of voxels that went into decoders were 2036, 2016, and 1433 for superficial, middle, and deep layer groups in FFA, and 1443, 1392, and 1151 for superficial, middle, and deep layer groups in LOC, respectively. To evaluate the potential bias due to this selection process choice of the “others” volumes, we also performed a permutation test by randomising correspondence between labels volumes one shuffle per target/”others” definition, which ended up with 100 subsets in total. For statistical analysis, we first performed one-way analysis of variance (ANOVA) for each comparison across layers with Bonferroni correction across labels and ROIs (14 labels x 2 ROIs). Only labels that showed significant differences were included in two-sample t-tests with Bonferroni correction across layers. Python 3.8.5 scipy 1.15.1 was used for both statistical analyses.

## Results

We confirmed the image distortion was mild around FFA, where it is typically very challenging to acquire reasonable signals (Fig. 2). For your inspection, browse the dataset (https://openneuro.org/datasets/ds003216).

**Figure 1.**
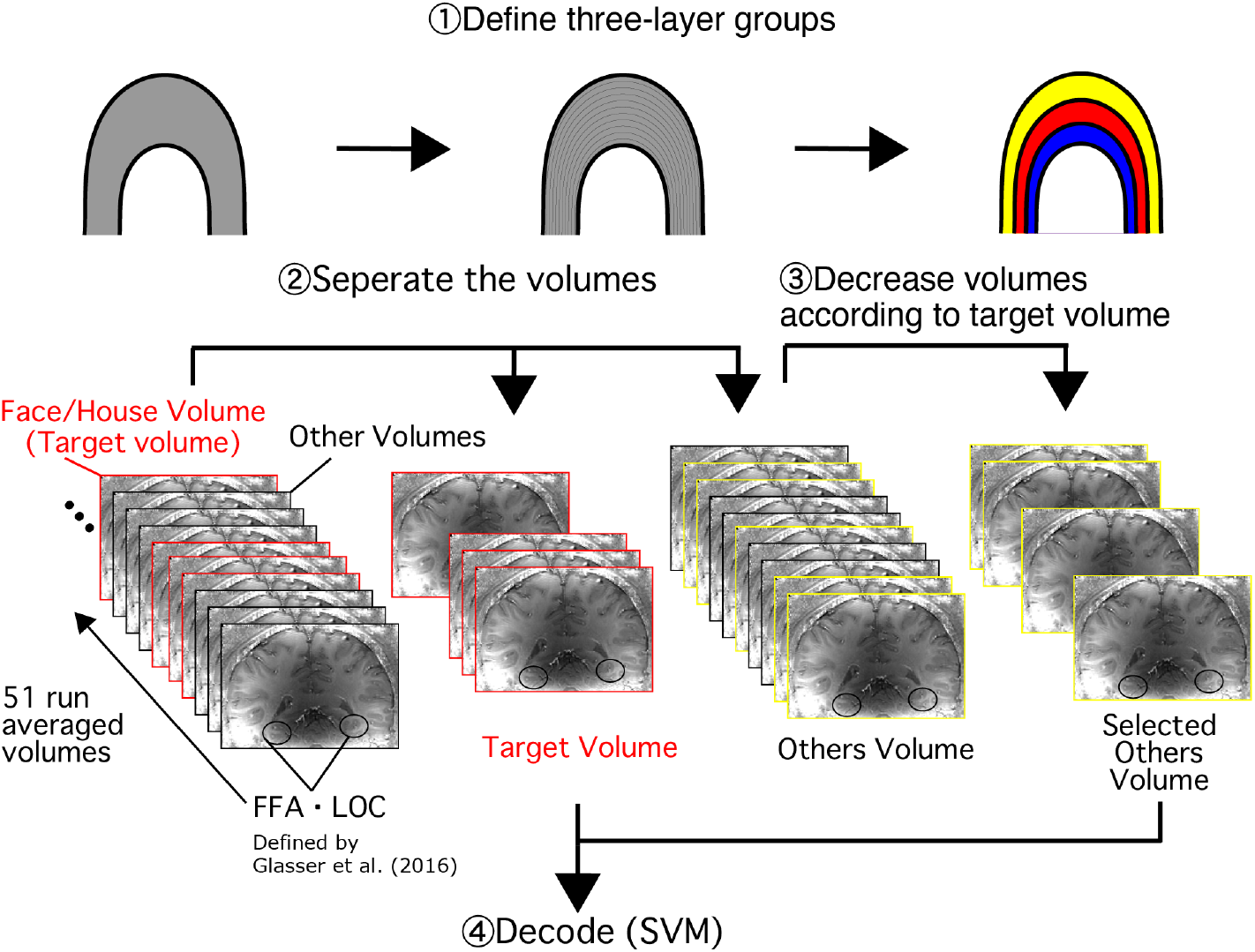
Analysis pipeline.

**Figure 2.**
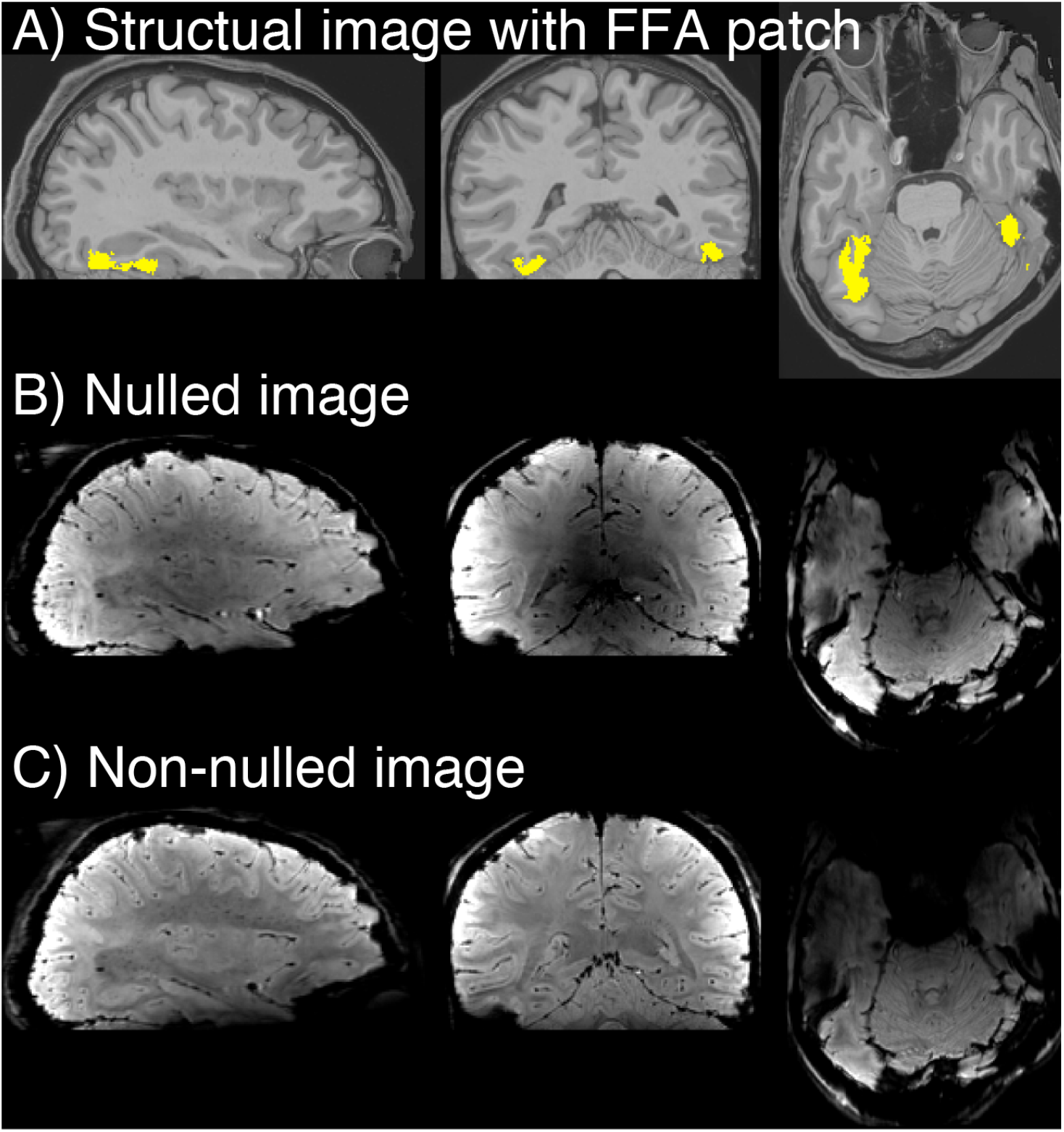
Image quality around the fusiform face area. Structural image with FFA patch in yellow (panel A), blood-nulled (VASO) image (panel B), and non-nulled (BOLD) image (panel C) are presented.

In VASO data, face-vs.-others and man-vs.-others decoding accuracy peaked in the middle layers in FFA with statistical significance (ANOVA: Bonferroni-corrected p < 0.05, pair-wise one-sided t-test superficial vs. middle layer: p < 0.05 (Bonferroni corrected), pair-wise one-sided t-test middle vs. deep layer: p < 0.05 (Bonferroni corrected)) (Fig. 3 & Fig. S1). However, the other decoding accuracies for any other object labels did not peak in the middle layers in FFA and did not show statistical significance in both superficial vs. middle layer and middle vs. deep layer. In LOC, none of the object labels showed a clear peak of decoding accuracy for the middle layers (Fig. S1). While VASO showed significantly different layer modulations (Fig. 3A), BOLD did not detect unique signal modulations across layers in either face-vs.-others or house-vs.-others decoders (Fig. 3B). For all other labels, BOLD did not detect unique signal modulation across layers, except man-vs.-others decoder (Fig. S2). This overall trend in BOLD might be due to expected venous drainage or high physiological noise.

**Figure 3.**
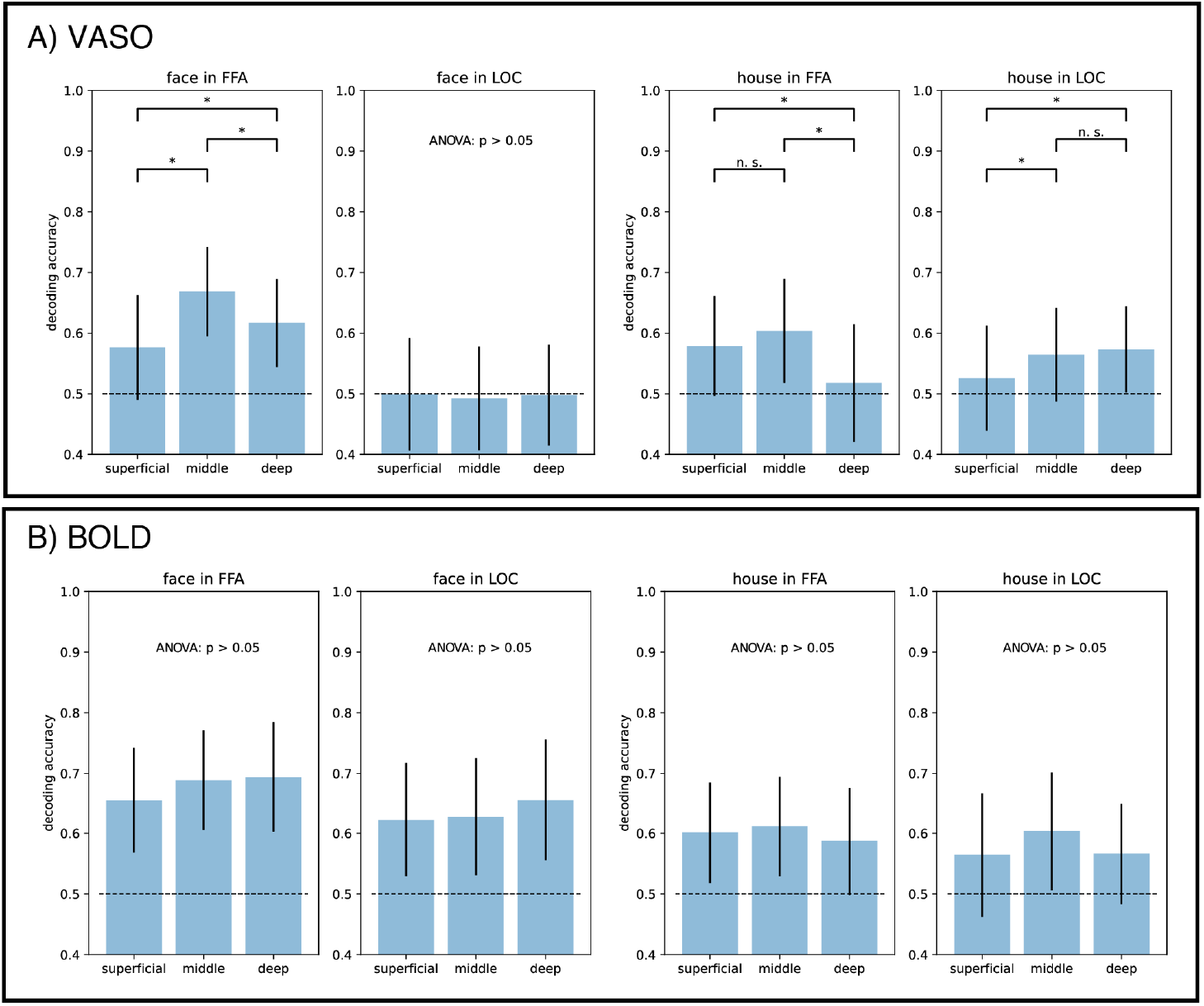
Decoding accuracy for each layer group. Panel A shows the decoding accuracy using VASO data. Panel B shows the decoding accuracy using BOLD data. Means and standard errors across 100 iterations of subgroup choices were plotted. Two-sample t-tests were performed and showed significance as star symbols (p < 0.05, Bonferroni corrected) in case one-way ANOVA showed significance (p < 0.05, Bonferroni corrected). Barplot, error bars, and statistical indications mean the same for all other figures below.

When permutation tests were performed, all the decoding accuracy became around the chance level and no significant peak for the middle layer was found in any labels for VASO and BOLD data (Fig. 4, Fig. S3, and Fig. S4). These results suggest that there was no critical information leakage due to the selection of training and test datasets. Namely, the decoders’ performance was based on statistical correspondence between fMRI data and labels, not based on random noise information.

**Figure 4.**
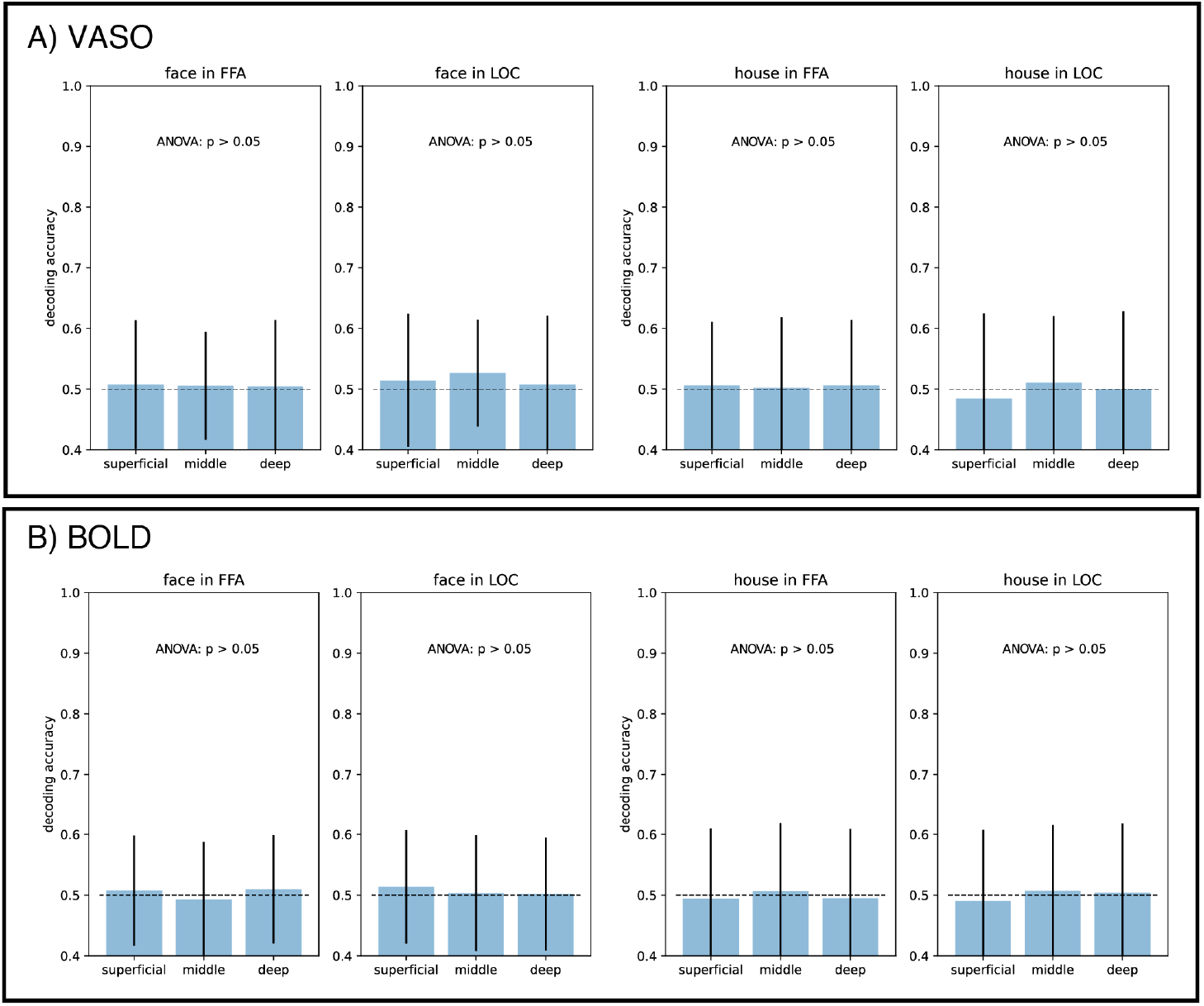
Decoding accuracy for each cortical layer with label permutations. Panel A shows the decoding accuracy using VASO data. Panel B shows the decoding accuracy using BOLD data.

## Discussion

In this study, we analysed an open whole-brain VASO layer fMRI dataset to investigate laminar-level representation in object-category recognition areas. Even though it is typically very challenging to acquire reliable signals from these lower areas of the brain, the image had decent contrast across tissues and did not show significant distortion (Fig. 2). Our results indicate that face information can be most accurately decoded in the middle layer in FFA, which corresponds to a face-representing subregion in the tested object-category areas. This suggests that face information in FFA is given by the feedforward neural signals during passive face image presentation.

### Locus of face information processing

We found that the face-vs.-others decoding accuracy in the middle layer in FFA was significantly higher than in other layers in VASO data. Most importantly, this signature was only depicted in this specific label and ROI combination but not in others. The middle-layer driven profile of the decoding performance suggests that FFA receives more feedforward signals rather than feedback signals during passive face image presentation. This could be further speculated that face information is already structured prior to FFA to some extent. Other studies (Nichols et al., 2010) showed that information about facial parts exists not only in FFA but also in OFA, which is believed as an area strongly related to partial processing of facial information earlier than FFA. They also suggest that complete face information is more pronouncedly represented in FFA. This agrees well with the results of the present study in the sense that face information has been partially processed before FFA, and hence FFA receives face information as a feedforward neural signature, and this partial information would be used to complete face representation as a whole in FFA.

However, face processing is very complicated and controversial, especially in terms of “where” the face information is structured. Bernstein and Yovel reviewed the past 20 years of literature (Bernstein & Yovel, 2015). They proposed that face processing involves two distinct neural pathways: static face processing, such as face identity and expression, and dynamic face processing, such as motion and expression. According to their study, information flows from OFA to FFA in the static face processing pathway, whereas it flows from OFA to posterior superior temporal sulcus (pSTS) in the dynamic face processing pathway. Our task, on the other hand, was to present face images in clips of movies, which included both facial expression and some motion, and the participant was not instructed to do any particular identification (no face identification task nor expression identification task). Hence, we cannot speculate that one of the other pathways was dominantly used in our paradigm. In this study, we analysed FFA to test the hypothesis of where the face information is structured. This means that it would also be worthwhile to investigate whether pSTS also has some information about the face, namely, how face image decoding accuracy would vary across cortical layers in pSTS. Also, Dowdle et al. (2021) comprehensively explored face-related areas with ambiguous tasks and showed feedback signatures in FFA. This discrepancy with the current study might come from the difference in the experimental tasks: our study used passive face image presentation, whereas Dowdle et al. (2021) used a task requiring active face detection in response to various ambiguous face stimuli. Further work should be done to comprehensively investigate whether the task engagement alters the face information pathway, feedforward or feedback signature, in the relevant brain regions.

Yet, as far as the authors’ knowledge, this is the first VASO layer fMRI work that investigated FFA in terms of face recognition pathway using MVPA. This paves the way to further investigate the face recognition pathway with ultra-high field fMRI in combination with MVPA.

### Training and test dataset sampling for decoding

In our analysis, due to the low SNR of the dataset, we first averaged all the runs and defined the training/test dataset in volume by volume. In this way, we were cautious about information leakage across volumes since we analyzed a single, averaged continuous run while splitting it into training and test trials, rather than using independent runs. However, our permutation test showed that all the decoding accuracies dropped to the chance level. This suggests that the decoding accuracy for the original, non-permuted test was not achieved by information leaked across training and test trials.

There would be another concern about sampling volumes with “others” labels since they were randomly chosen from the remaining volumes after picking the target volumes and removing all volumes contaminated by target labels. To evaluate the potential bias due to this selection process, we repeated the random sampling of the “others” volumes 100 times with replacement. As shown in Fig. 3, “face” and “man” labels in FFA had statistically significant accuracy for the middle layer, implying the existence of feedforward signatures in FFA.

However, BOLD showed decoding accuracy peaked in the middle layer for man-vs.-others classification in LOC (Fig. S2). This is not the result we expected from decoding results with other object labels and ROIs. We interpret this discrepancy as a result of the sampling procedure for the “man” label and the accompanying statistical bias. As mentioned above, we repeated the random sampling of the “others” volumes 100 times to mitigate the potential bias, given that the number of target volumes was much smaller than the corresponding “others” volumes for most labels. However, “man” and “sky” labels showed up far more frequently in the stimulus clips than other labels (63 volumes with the “man” label and 55 volumes with the “sky” label, whereas 19.3 volumes with a standard error of 2.3 for other labels). This restricted variations in the random sampling of the corresponding “others” volumes (ratios of “others” volumes actually chosen to all “others” volumes were 98.4 % for man and 80.9 % for sky, whereas 17.2 % with a standard error of 9.9% for other labels). Hence, if the statistical bias is introduced by using the fixed sampling of volumes, it cannot be mitigated by the iterative sampling for these two labels. This may explain the unexpected layer-dependent decoding pattern observed in the LOC for the “man” label in the BOLD signal.

### Precision scanning with one single participant

In this work, we analysed a single-subject deep-sampled dataset with over 10 two-hour sessions of 7T fMRI scanning. Typically, the lower parts of the cortex are challenging to acquire signals because the receiver RF-coil is located far from these areas, and there are stronger B0 inhomogeneities around them. In addition, this dataset was acquired by VASO, which has lower sensitivity than the “gold standard” fMRI contrast, GE-BOLD, even though VASO has higher specificity (Huber et al., 2019). This is why averaging all these runs was required.

Although only a single participant was scanned in this dataset, to our knowledge, this is the only publicly open dataset that allows us to analyze signals in the lower areas of the cortex with higher laminar specificity. Even though VASO has lower sensitivity compared to other contrasts, it is a quantitative contrast with higher spatial specificity. The characteristics indeed helped us to see laminar-wise differences in signals, especially in combination with MVPA, which enables us to access information contents in each layer that mere univariate analysis cannot retrieve (see Fig. 3).

However, it might be risky to overinterpret the results from a single participant. Typical neuroimaging studies include far more participants, yet with far less data per participant, to assess group-level consistency or to examine individual variability while minimizing the risk of spurious findings. In this study, we have used a deep-sampled dataset, in which a single participant was scanned over 10 sessions. The amount of data is equivalent to 10 participants with a single session per participant, except that it is from the same brain. Moreover, to make sure our result was not a spurious finding, other labels were also analysed and compared with the target (“face”). The results showed that the feedforward signature was uniquely found for “face-vs.-others” decoding in FFA, except for one case in BOLD data (see above for details). Also, we tested the permutation of labels to ensure the effect of the autocorrelation did not affect our results. All of these analyses suggest that the results found in this study are neuroscientifically meaningful, but not spurious findings.

Although we still need to be cautious about individual variability, the visual organization we investigate in this study is consistent across individuals (Naselaris et al., 2021). Hence, it seems unlikely that only this participant showed this trend in the data and other participants did not.

This is why investigating a single brain is also useful, especially in the brain areas where we do not expect major individual variability. Yet, it would be ideal to test a few more participants to make sure it is not due to individual variability.

For the dataset we used in this study, we tried to push the limit of layer fMRI to acquire decent signals from a lower part of the brain. To realize this, we performed many piloting sessions, more than 30 scanning hours, to find the best MR sequence for this particular paradigm. We hope this work will be a piece of evidence for VASO’s usability and will showcase the unique insight and additional information that VASO can provide.

### Choice of control ROI

In this study, we investigated the MR signal in FFA and LOC as examples of object representation areas. Parahippocampal place area (PPA) was not included in our analysis because the movie stimulus used in this study consistently contained scene images. Since PPA is known to respond strongly to scene images (Epstein et al., 1999), we anticipated continuous activation of this region throughout the paradigm. Consequently, it would not have been possible to assess label-dependent differences in brain activity patterns within PPA. For this reason, PPA was deemed unsuitable for comparison with FFA and was therefore excluded from further analysis.

### Comparison to BOLD signal

In this study, we compared layer-dependent decoding accuracies between VASO and BOLD data. We should still note that BOLD data in this dataset were acquired for the sake of BOLD correction for VASO data, namely, the image acquisition parameters were not optimized to acquire BOLD data per se. However, decoding accuracies for BOLD data were marked as high as, or even higher than those for VASO data in most labels in both regions of interest. On the other hand, the VASO data showed a feedforward signature in decoding accuracies for face-vs.-others and man-vs.-others in FFA, but the BOLD data did not. This is consistent with the previous literature that the BOLD signal has higher sensitivity but the VASO signal has higher specificity (e.g., Huber et al., 2019). Therefore, this is the additional value of using VASO even if its SNR is much less than BOLD.

### Another limitation in the dataset

It is known that scanning the lower brain area at high spatial resolution is very challenging, as high spatial resolution scanning introduces a low spatial frequency EPI artifact, also known as the Fizzy Ripple artifact (Huber et al., 2025). In the future, we might overcome this low SNR issue and eliminate the need for deep sampling by utilizing methods such as complex-valued averaging of dual-polarity EPI to reduce Fizzy Ripple artifact (see Fig. 7 in Huber et al., 2025).

## Conclusions

In this study, we investigated face information representation across cortical layers in the visual object category-selective areas, fusiform face area (FFA) and lateral occipital cortex (LOC), using multi-voxel pattern analysis (MVPA). We used an open dataset from our previous study (Koiso et al., 2023) to examine whether face information processing occurs before FFA—that is, whether face information is represented in FFA as a feedforward signature. Our result supported our hypothesis by showing a feedforward signature in FFA for face information, but not for other object images. This pattern was not observed in BOLD data. These findings demonstrate that VASO layer fMRI can provide complementary insights to conventional BOLD layer fMRI, particularly when combined with MVPA.

## Data and Code Availability

Dataset: https://openneuro.org/datasets/ds003216

## Author Contributions

Conceptualization: K.K., K.A., and Y.M.; Data collection: K.K. and R.H.; Data analysis: K.K.; Visualization: K.K.; Writing—original draft: K.K. and Y.M.; Writing—review & editing: K.K., K.A., R.H., and Y.M.; Funding acquisition: R.H. and Y.M.

## Funding

NWO VENI project 016.Veni.198.032.

JSPS KAKENHI (25H01138, 20H00600, 18KK0311).

## Reference

Bernstein, M., & Yovel, G. (2015). Two neural pathways of face processing: A critical evaluation of current models. Neuroscience & Biobehavioral Reviews, 55, 536–546. 10.1016/j.neubiorev.2015.06.010

Carricarte, T., Iamshchinina, P., Trampel, R., Chaimow, D., Weiskopf, N., & Cichy, R. M. (2024). Laminar dissociation of feedforward and feedback in high-level ventral visual cortex during imagery and perception. iScience, 27(7), 110229. 10.1016/j.isci.2024.110229

Dowdle, L., Ghose, G., Moeller, S., Ugurbil, K., Yacoub, E., & Vizioli, L. (2022). Characterizing top-down microcircuitry of complex human behavior across different levels of the visual hierarchy. 10.1101/2022.12.03.518973

Epstein, R., Harris, A., Stanley, D., & Kanwisher, N. (1999). The Parahippocampal Place Area. Neuron, 23(1), 115–125. 10.1016/s0896-6273(00)80758-8

Glasser, M. F., Coalson, T. S., Robinson, E. C., Hacker, C. D., Harwell, J., Yacoub, E., Ugurbil, K., Andersson, J., Beckmann, C. F., Jenkinson, M., Smith, S. M., & Van Essen, D. C. (2016). A multi-modal parcellation of human cerebral cortex. Nature, 536(7615), 171–178. 10.1038/nature18933

Grill-Spector, K., Weiner, K. S., Kay, K., & Gomez, J. (2017). The Functional Neuroanatomy of Human Face Perception. Annual Review of Vision Science, 3(1), 167–196. 10.1146/annurev-vision-102016-061214

Haenelt, D., Chaimow, D., Nasr, S., Weiskopf, N., & Trampel, R. (2023). Decoding of columnar-level organization across cortical depth using BOLD- and CBV-fMRI at 7 T [Preprint]. Neuroscience. 10.1101/2023.09.28.560016

Huber, L., Ivanov, D., Krieger, S. N., Streicher, M. N., Mildner, T., Poser, B. A., Möller, H. E., & Turner, R. (2014). Slab-selective, BOLD-corrected VASO at 7 Tesla provides measures of cerebral blood volume reactivity with high signal-to-noise ratio: SS-SI-VASO MeasuresChanges of CBV in Brain. Magnetic Resonance in Medicine, 72(1), 137–148. 10.1002/mrm.24916

Huber, L., Stirnberg, R., Morgan, A. T., Feinberg, D. A., Ehses, P., Knudsen, L., Gulban, O. F., Koiso, K., Gephart, I., Swegle, S., Wardle, S. G., Persichetti, A. S., Beckett, A. J. S., Stöcker, T., Boulant, N., Poser, B. A., & Bandettini, P. A. (2025). Short-term gradient imperfections in high-resolution EPI lead to Fuzzy Ripple artifacts. Magnetic Resonance in Medicine, 94(2), 571–587. 10.1002/mrm.30489

Huber, L., Uludağ, K., & Möller, H. E. (2019). Non-BOLD contrast for laminar fMRI in humans: CBF, CBV, and CMRO2. NeuroImage, 197, 742–760. 10.1016/j.neuroimage.2017.07.041

Iamshchinina, P., Haenelt, D., Trampel, R., Weiskopf, N., Kaiser, D., & Cichy, R. M. (2021). Benchmarking GE-BOLD, SE-BOLD, and SS-SI-VASO sequences for depth-dependent separation of feedforward and feedback signals in high-field MRI. 10.1101/2021.12.10.472064

Kanwisher, N., McDermott, J., & Chun, M. M. (1997). The Fusiform Face Area: A Module in Human Extrastriate Cortex Specialized for Face Perception. The Journal of Neuroscience, 17(11), 4302–4311.10.1523/JNEUROSCI.17-11-04302.1997

Koiso, K., Müller, A. K., Akamatsu, K., Dresbach, S., Wiggins, C. J., Gulban, O. F., Goebel, R., Miyawaki, Y., Poser, B. A., & Huber, L. (2023). Acquisition and processing methods of whole-brain layer-fMRI VASO and BOLD: The Kenshu dataset. Aperture Neuro, 3. 10.52294/001c.87961

Mekete, N., & Vu, A. (2017). Encoding and decoding semantic information of natural movies from 7T human brain activity provided by the Human Connectome Project. ISMRM, 5369. Moerel, M., De Martino, F., Kemper, V.G., Schmitter, S., Vu, A.T., Uğurbil, K., Formisano, E., & Yacoub, E. (2018). Sensitivity and specificity considerations for fMRI encoding, decoding, and mapping of auditory cortex at ultra-high field. NeuroImage, 164, 18–31. 10.1016/j.neuroimage.2017.03.063

Muckli, L., De Martino, F., Vizioli, L., Petro, L. S., Smith, F. W., Ugurbil, K., Goebel, R., & Yacoub, E. (2015). Contextual Feedback to Superficial Layers of V1. Current Biology, 25(20), 2690–2695. 10.1016/j.cub.2015.08.057

Naselaris, T., Allen, E., & Kay, K. (2021). Extensive sampling for complete models of individual brains. Current Opinion in Behavioral Sciences, 40, 45–51. 10.1016/j.cobeha.2020.12.008

Nichols, D. F., Betts, L. R., & Wilson, H. R. (2010). Decoding of faces and face components in face-sensitive human visual cortex. Frontiers in Psychology. 10.3389/fpsyg.2010.00028

Poser, B. A., Ivanov, D., Kemper, VG, Kannengiesser, SA, Uludag, K, & Barth, M. (2013). CAIPIRINHA-accelerated 3D EPI for high temporal and / or spatial resolution EPI acquisitions. Esmrmb.

Sellal, F. (2022). Anatomical and neurophysiological basis of face recognition. Revue Neurologique, 178(7), 649–653. 10.1016/j.neurol.2021.11.002

Vizioli, L., De Martino, F., Petro, L. S., Kersten, D., Ugurbil, K., Yacoub, E., & Muckli, L. (2020). Multivoxel Pattern of Blood Oxygen Level Dependent Activity can be sensitive to stimulus specific fine scale responses. Scientific Reports, 10(1). 10.1038/s41598-020-64044-x

