## Supplemental Figure 1-4 for "Feedforward representation of face information in the fusiform face area revealed by VASO 7T layer fMRI"

### Supplementary Material

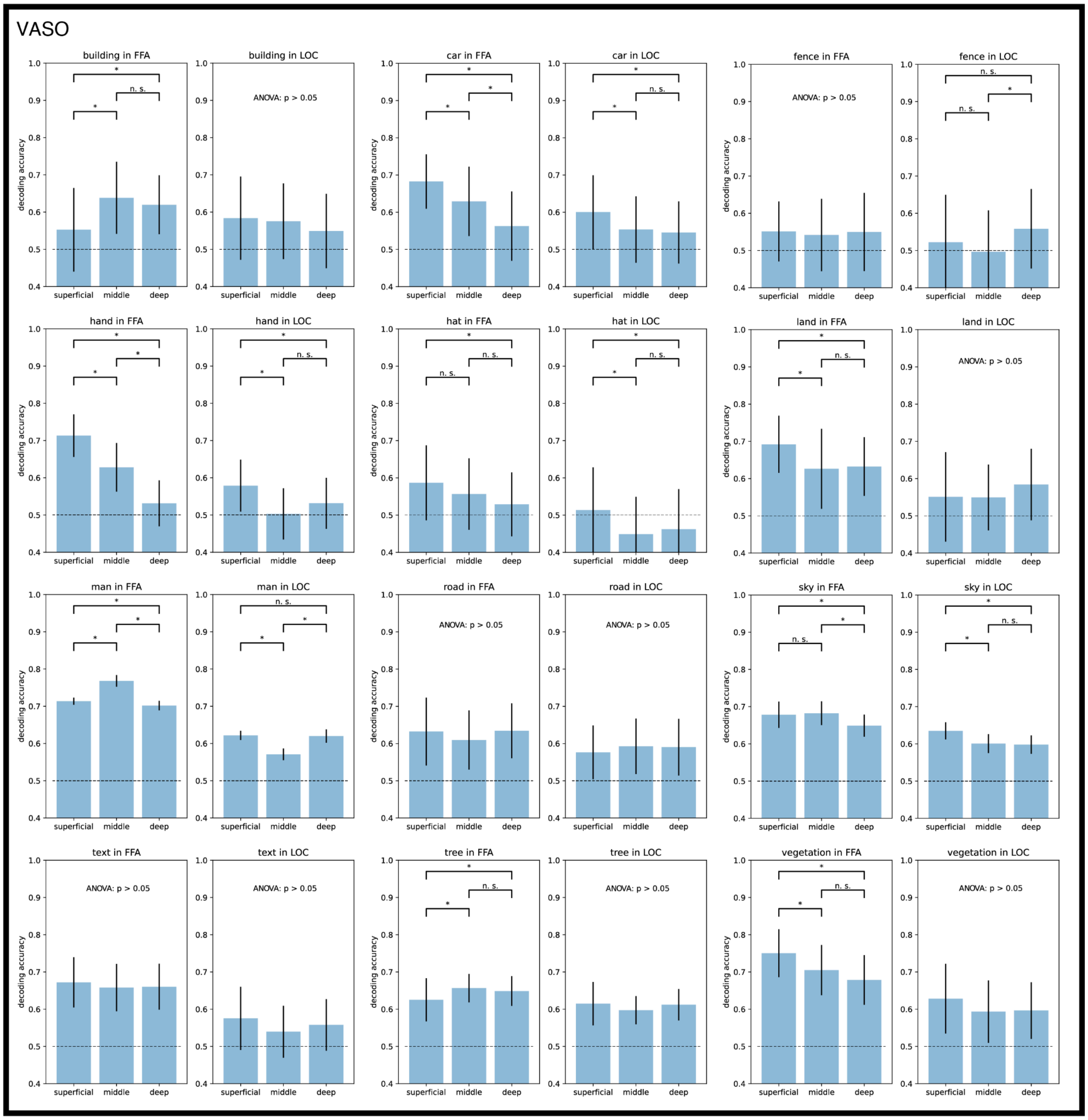


*Figure S1. Decoding accuracy for each cortical layer group in VASO.*
*Means and standard errors across 100 iterations of subgroup choices were plotted. Two-sample t-tests were performed and showed significance as star symbols ($p < 0.05$, Bonferroni corrected) in case one-way ANOVA showed significance ($p < 0.05$, Bonferroni corrected). Barplot, error bars, and statistical indications mean the same for all other figures below.*

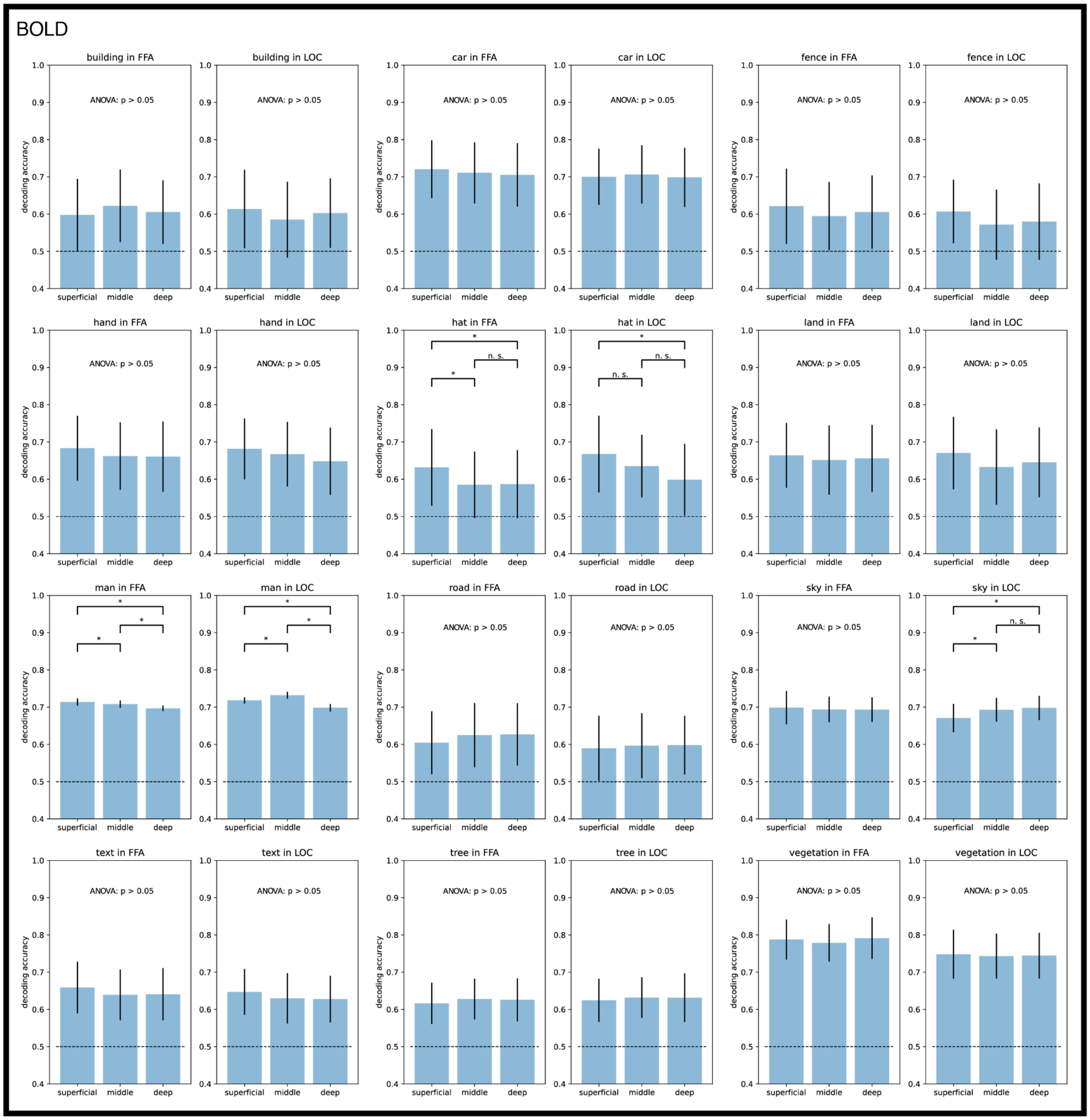


*Figure S2. Decoding accuracy for each cortical layer group in BOLD.*

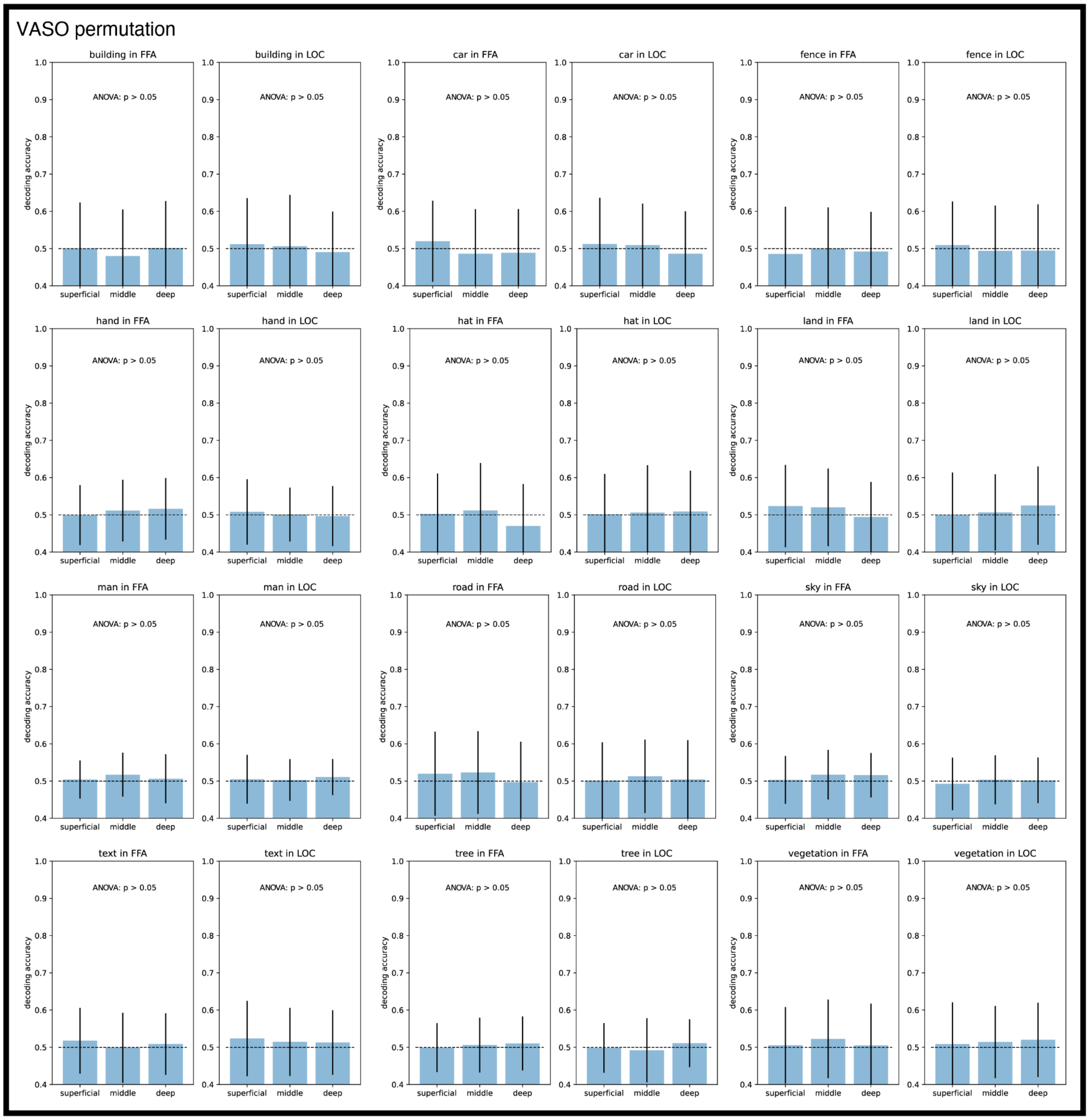


*Figure S3. Decoding accuracy for each cortical layer group in VASO with label permutations.*

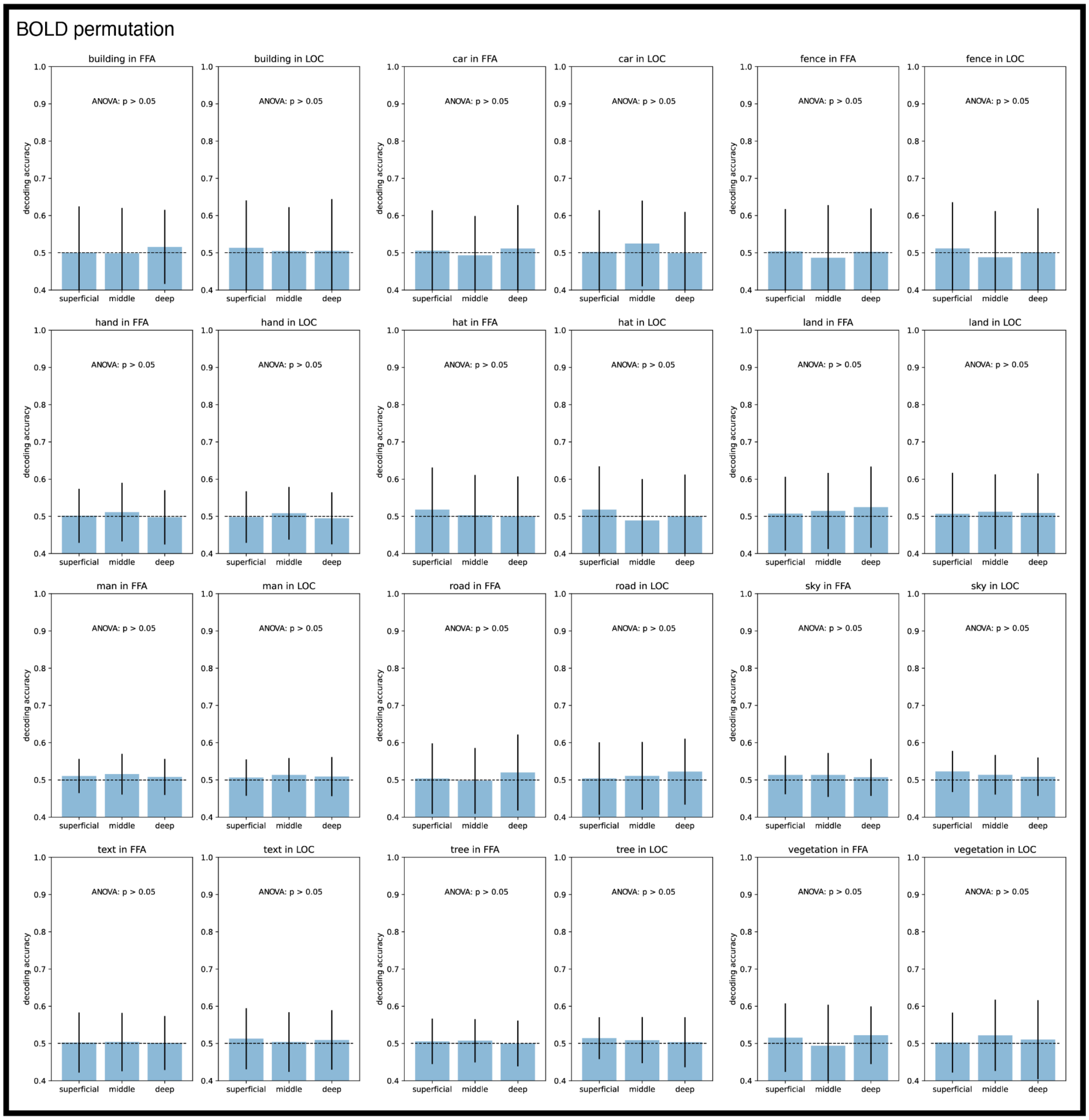


*Figure S4. Decoding accuracy for each cortical layer group in BOLD with label permutations.*
